# Evaluation of the performance of SARS-CoV-2 serological tools and their positioning in COVID-19 diagnostic strategies

**DOI:** 10.1101/2020.06.16.156166

**Authors:** Aurelie Velay, Floriane Gallais, Ilies Benotmane, Marie Josée Wendling, François Danion, Olivier Collange, Jérôme De Sèze, Catherine Schmidt-Mutter, Francis Schneider, Pascal Bilbault, Ferhat Meziani, Samira Fafi-Kremer

**Affiliations:** Laboratoire de Virologie, Hôpitaux Universitaires de Strasbourg, Strasbourg, France; INSERM, UMR_S1109, LabEx TRANSPLANTEX, Centre de Recherche d’Immunologie et d’Hématologie, Faculté de Médecine, Fédération Hospitalo-Universitaire (FHU) OMICARE, Fédération de Médecine Translationnelle de Strasbourg (FMTS), Université de Strasbourg, Strasbourg, France; Service des Maladies Infectieuses et Tropicales, Hôpitaux Universitaires de Strasbourg, Strasbourg, France; Département d’Anesthésie et Réanimation Chirurgicale, Hôpitaux Universitaires de Strasbourg, Strasbourg, France; Centre d’Investigation Clinique INSERM CIC – 1434, Hôpitaux Universitaires de Strasbourg, Strasbourg, France; Département de Médecine Intensive-Réanimation, Hôpitaux Universitaires de Strasbourg, Strasbourg, France; Département des Urgences Médicales, Hôpitaux Universitaires de Strasbourg, Strasbourg, France; Service de Médecine Intensive-Réanimation, Hôpitaux Universitaires de Strasbourg, Strasbourg, France; INSERM, UMR 1260, Regenerative Nanomedicine (RNM), FMTS, Strasbourg, France

## Abstract

Rapid and accurate diagnosis is crucial for successful outbreak containment. During the current coronavirus disease 2019 (COVID-19) public health emergency, the gold standard for severe acute respiratory syndrome coronavirus 2 (SARS-CoV-2) infection diagnosis is the detection of viral RNA by reverse transcription (RT)-PCR. Additional diagnostic methods enabling the detection of current or past SARS-CoV-2 infection would be highly beneficial to ensure the timely diagnosis of all infected and recovered patients. Here, we investigated several serological tools, i.e., two immunochromatographic lateral flow assays (LFA-1 (Biosynex COVID-19 BSS) and LFA-2 (COVID-19 Sign IgM/IgG)) and two enzyme-linked immunosorbent assays (ELISAs) detecting IgA (ELISA-1 Euroimmun), IgM (ELISA-2 EDI) and/or IgG (ELISA-1 and ELISA-2) based on well-characterized panels of serum samples from patients and healthcare workers with PCR-confirmed COVID-19 and from SARS-CoV-2-negative patients. A total of 272 serum samples were used, including 62 serum samples from hospitalized patients (panel 1 and panel 3), 143 serum samples from healthcare workers (panel 2) diagnosed with COVID-19 and 67 serum samples from negative controls. Diagnostic performances of each assay were assessed according to days after symptom onset (dso) and the antigenic format used by manufacturers. We found overall sensitivities ranging from 69% to 93% on panels 1 and 2 and specificities ranging from 83% to 98%. The clinical sensitivity varied greatly according to the panel tested and the dso. The assays we tested showed poor mutual agreement. A thorough selection of serological assays for the detection of ongoing or past infections is advisable.

## INTRODUCTION

A novel coronavirus named severe acute respiratory syndrome coronavirus 2 (SARS-CoV-2) causing coronavirus disease 2019 (COVID-19) has emerged as a major healthcare threat (1). At the beginning of the pandemic, the main healthcare objective was to stop the spread of the virus. A key aspect to achieve this goal was to ensure early and accurate infection diagnosis and appropriate quarantine for infected people. The gold standard for identifying SARS-CoV-2 infection relies on the detection of viral RNA by reverse transcription (RT-) polymerase chain reaction (PCR)-based techniques. However, the large-scale routine implementation of this approach has been hampered by its time-consuming nature (most often 4–6 hours) and shortages of materials. Moreover, the presence of sufficient amounts of the viral genome at the site of sample collection is a prerequisite to allow genome detection. Missing the time window of active viral replication or low-quality sampling can lead to false-negative results, which would allow infected patients to spread the virus to their relatives and working environment. In such conditions, additional diagnostic methods would be highly beneficial to ensure timely diagnosis of all infected and recovered patients. Combining RT-PCR with the screening of the onset and strength of the humoral response against SARS-CoV-2 could enhance diagnostic sensitivity and accuracy. There are now several studies describing the kinetics of anti-SARS-CoV-2 IgM and IgG detection using laboratory ELISA tests, most reporting that IgM is detectable as early as 5-14 days after the first clinical symptoms (2–7). At this stage of the pandemic, many countries are now questioning how to prepare and manage the easing of lockdown. Serological tools have an important place in establishing such strategies. Validated serological assays are crucial for patient contact tracing and epidemiological studies. Several formats of serological methods are beginning to be marketed, i.e., lateral flow assays (LFAs) and enzyme-linked immunosorbent assays (ELISAs) detecting IgA, IgM and/or IgG. Data about the analytical and clinical performances of these devices are still lacking, as well as their indication in the diagnosis of SARS-CoV-2 infection. In this context, we evaluated the diagnostic performances of two LFAs and two commercial ELISA kits detecting IgM, IgA and IgG based on well-characterized panels of serum samples from PCR-confirmed COVID-19 patients and healthcare workers and from SARS-CoV-2-negative patients. Diagnostic performances of each assay were assessed according to days after symptom onset (dso) and the antigenic format used by manufacturers. This evaluation led us to propose a decisional diagnostic algorithm based on serology, which may be applicable in future seroprevalence studies.

## MATERIALS AND METHODS

### Patients and serum samples/Study design

The study design is summarized in Figure 1. A total of 272 serum samples were used, including 62 serum samples from hospitalized patients (30 of the 62 in panel 1 and 50 of the 62 in panel 3); 143 serum samples from healthcare workers (panel 2) diagnosed with COVID-19 at Strasbourg University Hospital (Strasbourg, France), recruited in April 2020; and 67 serum samples from negative controls. All sera of panels 1 and 2 were tested with two LFAs and two IgG ELISAs (Fig. 1). Fifty serum samples (panel 3) from infected patients collected from 1 to 14 dso were tested by IgA and IgM ELISA. Patient characteristics (the date and type of presenting symptoms) were collected for each panel (Table 1). Laboratory detection of SARS-CoV-2 was performed by RT-PCR testing of nasopharyngeal swab specimens according to current guidelines (Institut Pasteur, Paris, France; WHO technical guidance). This assay targets two regions of the viral RNA-dependent RNA polymerase (RdRp) gene, with a threshold limit of detection of 10 copies per reaction. Serum samples were collected at a median of 9 dso (range, 1-28 dso) for panel 1, 24 dso (range, 15-39 dso) for panel 2, and 7 dso (range, 0-14 dso) for panel 3. Serum samples from 40 patients collected before the COVID-19 pandemic onset (from March to November 2019) were selected as negative controls to determine clinical specificity. Another 27 serum samples were used to study cross-reactivity, including 20 samples from patients infected with four other human coronaviruses two to three months before sampling (HCoV-229E, HCoV-HKU1, HCoV-NL63, and HCoV-OC43), two from patients previously infected with influenza A virus, one from a patient previously infected with human rhinovirus, two containing rheumatoid factor, and two positive for antinuclear antibodies. All these negative controls were tested with all evaluated assays. Additionally, nine lots of intravenous immunoglobulins and one pool of six solvent detergent fresh-frozen plasma bags from healthy donors obtained from the French blood bank were tested. Ethical approval was granted by the local institutional review board (CE-2020-34). All patients provided written informed consent.

**Figure 1.**
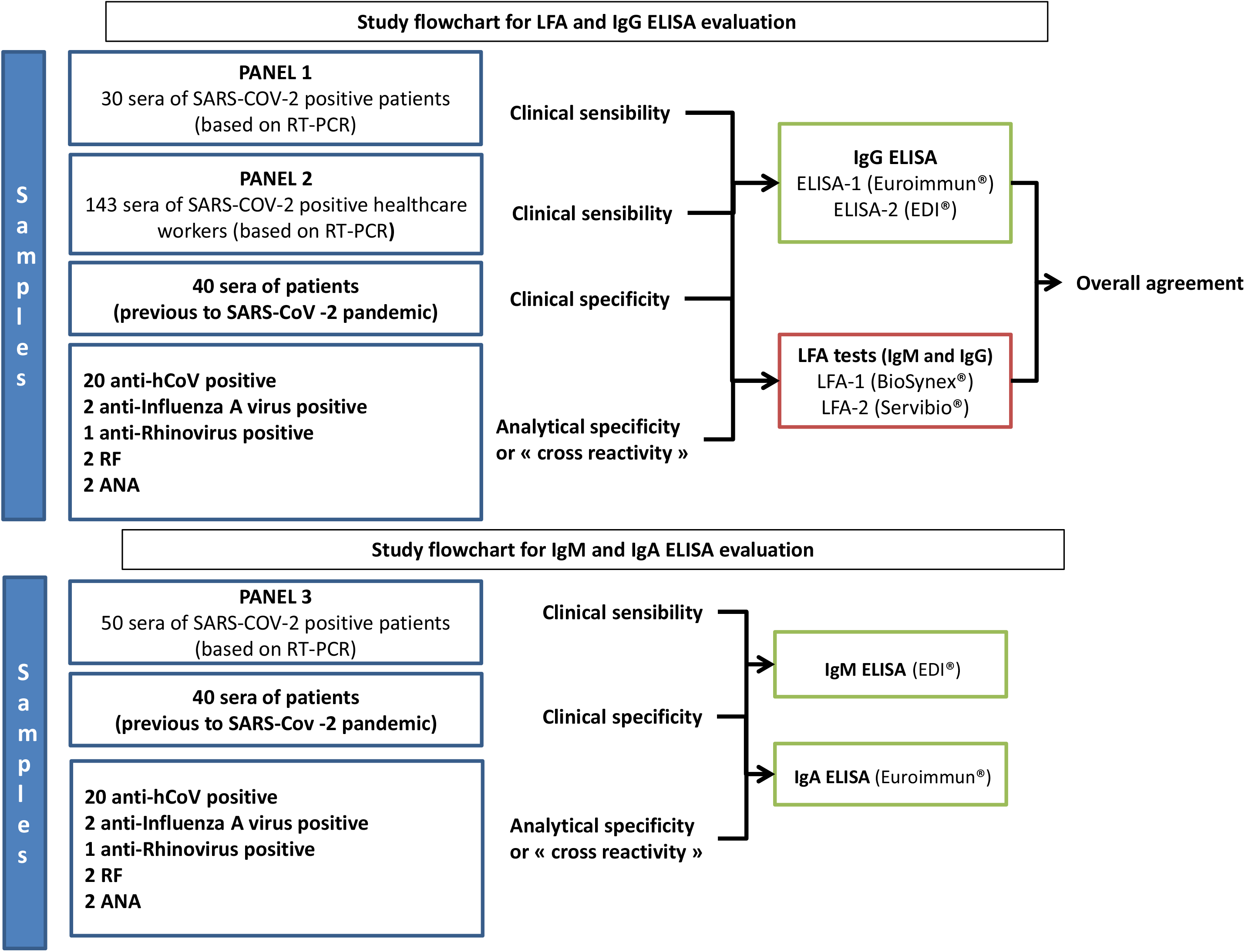
Study flowchart for LFA and ELISA evaluation. Panel 1 and panel 2 were used to determine the clinical sensitivity of the LFA and IgG ELISA. Panel 3 was used to determine the clinical sensitivity of the IgA and IgM ELISA. RF corresponds to samples containing rheumatoid factor, and ANA refers to samples containing antinuclear antibodies.

**Table 1:** Patient characteristics

|  | COVID-19<br>patients<br>(panel 1 and panel 3) | COVID-19<br>healthcare workers<br>(panel 2) | Total |
| --- | --- | --- | --- |
| Number of patients | 55 | 143 | 198 |
| Median age (years)<br>[range] | 68<br>[34-93] | 32<br>[21-62] | 43<br>[21-93] |
| Sex (female/male) | 17/38 | 96/47 | 113/85 |
| Median dso at RT-PCR analysis<br>[range] | 3<br>[0-13] | 2<br>[0-11] | 2<br>[0-13] |
| Median dso at serum collection<br>[range] | 8<br>[0-28] | 24<br>[15-39] | 22<br>[0-39] |
| Hospitalized in ICU | 23 | NA | NA |
| Hospitalized without ICU admission | 33 | NA | NA |

dso: Days after symptom onset
ICU: Intensive care unit
NA: Not applicable

Samples analyzed within 7 days were stored at 4°C. The other samples were stored at −20°C with only a single freeze-thaw cycle.

### Immunochromatographic lateral flow assays (LFAs)

We evaluated two commercial CE-marked LFAs: (i) LFA-1: Biosynex COVID-19 BSS (Biosynex, Switzerland, Fribourg) and (ii) LFA-2: COVID-19 Sign IgM/IgG (Servibio/VEDALAB, France, Alençon). Technical characteristics of the assays are summarized in the Supplementary data (Table S1). Both were tested according to the manufacturer’s instructions. Briefly, for each test, 10 μL of serum sample and two drops of buffer were added. The strip was placed flat at room temperature for 10 minutes, and then the results were scored according to the sample and control line intensity only for the tests validated by the appearance of the control line. Interpretation was performed by two independent readers using the standardized intensity scoring system that was established previously. The absence of the sample line was scored as 0 (negative), whereas a visible sample line was classified as positive, and the results were scored as follows: a weak line as 1, a clear visible line with an intensity lower than that of the control line as 2, a clear visible line with an intensity similar to that of the control line as 3, and a clear visible line with an intensity higher than that of the control line as 4.

### Enzyme-linked Immunosorbent Assay (IgA, IgM and IgG)

The following ELISA diagnostic kits were used for the detection of anti-SARS-CoV-2 IgA, IgM and IgG antibodies according to the manufacturer’s instructions: (1) ELISA-1: ELISA anti-SARS-CoV-2 IgA and IgG (Euroimmun, Lübeck, Germany) and (2) ELISA-2: EDI™ novel coronavirus COVID-19 IgM and IgG (Epitope Diagnostics, San Diego, CA, USA). Technical characteristics of the assays are summarized in the Supplementary data (Table S1). The assessed ELISA kits used as their antigenic source full-length recombinant nucleocapsid protein and the recombinant S1 domain of the spike protein for IgA and IgG in ELISA-1 and for IgM and IgG in ELISA-2, respectively. In brief, the optical density (OD) of the samples and calibrators was detected at 450 nm. Cutoffs for IgG detection were calculated according to the manufacturer’s instructions. ELISA-1 results were expressed as a ratio, and a ratio greater than 1.1 was considered positive. For ELISA-2, values greater than the cutoff were considered positive. To allow correlation of the results, the results for IgG ELISA-2 were also expressed as a ratio (OD sample/OD cutoff).

### Statistical analysis

Clinical sensitivity was determined on samples from SARS-CoV-2 RT-PCR-positive patients and healthcare workers (inclusion criterion). Percentages of IgA, IgM and IgG detection were calculated and compared among all evaluated serological devices according to the dso category in panel 1 (i.e., 1 to 7, 8 to 14, and more than 14), in panel 2 (i.e., 15 to 21, 22 to 28, and more than 28), and in panel 3 (i.e., 0 to 3, 4 to 8, and 9 to 14). For both LFAs, overall positivity was also evaluated based on positive results for the IgM or the IgG test line. Clinical specificity was calculated using the serum samples from 40 patients collected before the COVID-19 pandemic onset (from March to November 2019). Agreement among kits was determined for IgM and IgG parameters using Fleiss’ kappa (overall agreement) and Cohen’s kappa (agreement between pairs). A kappa value > 0.80 was deemed satisfactory. The diagnostic performances were estimated by comparing the combined IgM and IgG results according to SARS-CoV-2 infection status for each sample. Performance was considered satisfactory if the diagnostic accuracy exceeded 90%. Analyses were conducted using GraphPad (San Diego, CA, USA) Prism 6 software.

## RESULTS

### Study population

The general characteristics of the COVID-19 study participants are presented in Table 1. We collected serum samples from a total of 198 patients, including 85 men. Ages ranged from 21 to 93 years, with a median of 43. Serum samples were divided into several panels for evaluation, i.e., panels 1 and 3 correspond to COVID-19 patients, and panel 2 corresponds to COVID-19 healthcare workers. Among COVID-19 patients, the median age was 68 (range: 34-93), and the median age was 32 (range: 21-62) among COVID-19 healthcare workers.

### LFA and ELISA clinical performances

Clinical sensitivity and specificity The clinical sensitivity evaluated on 171 serum samples from COVID-19 patients (panel 1 and panel 2, excluding the second serum sample in repeatedly sampled patients) varied greatly between the two LFAs tested, especially for IgM, which was found in 83% and 30% of samples, respectively. A higher percentage of IgM detection (90%) was observed between 15 and 21 dso for LFA-1 (Fig. S1). The sensitivity was similar for IgG between the devices, with 68% of samples detected positive using LFA-1 and 65% detected positive using LFA-2. The maximal detection rate for IgG was observed 28 dso, with 88% and 80% for LFA-1 and LFA-2, respectively (Fig. S2). Combining IgM and IgG detection led to an overall sensitivity of 93% using LFA-1 but only 69% using LFA-2.

The clinical sensitivity estimated for IgG detection with ELISA-1 and ELISA-2 on the same 171 serum samples was 84% and 74%, respectively (Fig. 2). Both ELISA kits were more sensitive than the LFA devices for IgG detection between 22 and 28 dso. For this period, the sensitivity for IgG detection for ELISA-1 reached 96%.

**Figure 2.**
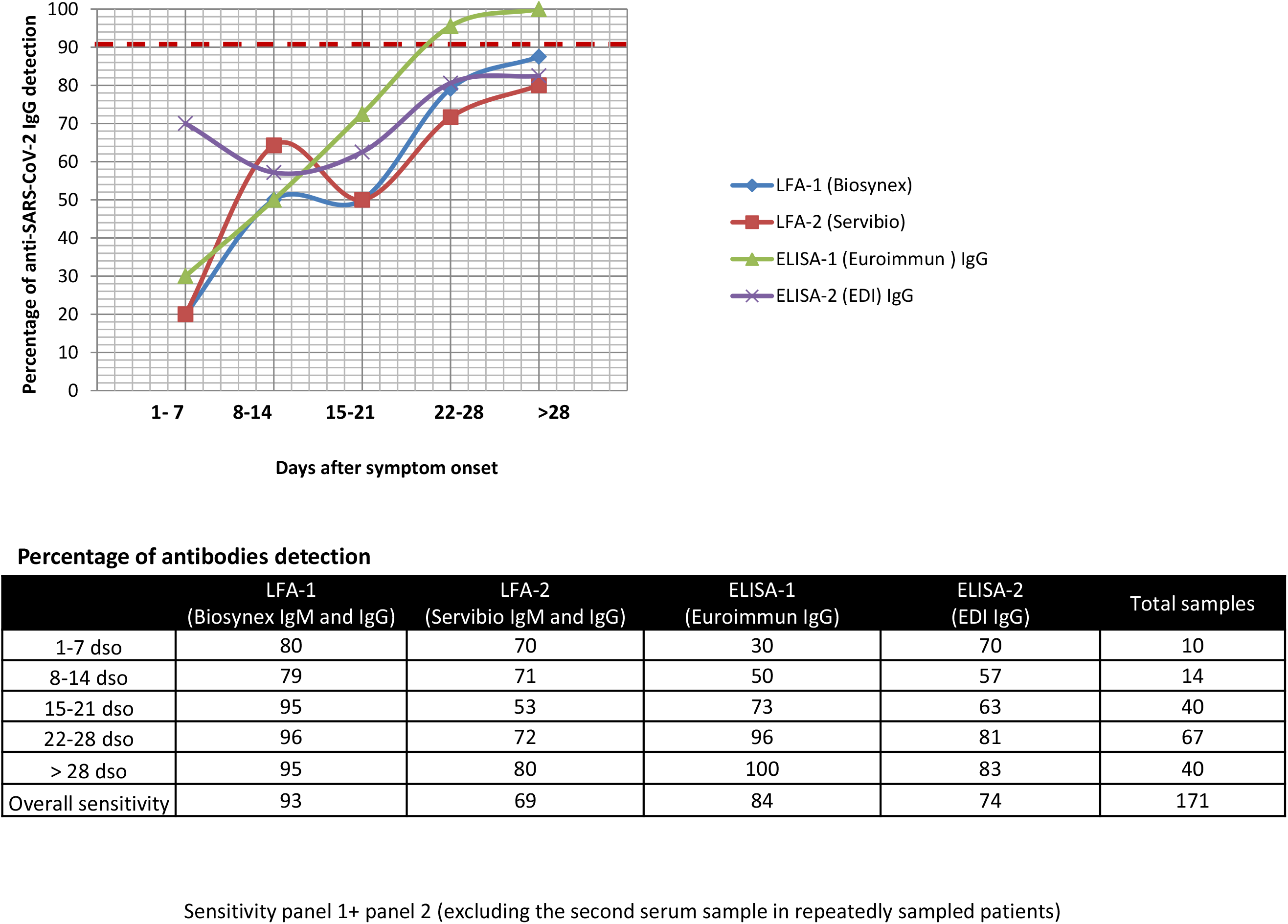
Rates of positivity for virus-specific antibodies measured by LFA (both IgG and IgM) and ELISA (IgG) versus time (in days) after the date of symptom onset from 171 COVID-19 hospitalized patients and healthcare workers.

Clinical sensitivity was also evaluated in each panel separately, given that specimens were sampled earlier after symptom onset in panel 1 than in panel 2. In panel 1, IgM was detected in 61% and 39% of sera using LFA-1 and LFA-2, respectively (Fig. S3). The optimum IgM detection rate was observed earlier with LFA-1 (80% of cases from 1 to 7 dso) than with LFA-2 (50% of cases only 8 dso). In this panel, the percentage of IgG detection ranged from 46% (LFA-1 and IgG ELISA-1) to 64% (IgG ELISA-2) (Fig. S4). The optimum rate of IgG detection was observed for all assays 14 dso, with rates ranging from 75% (LFA-2, IgG ELISA-1 and IgG ELISA-2) to 100% (LFA-1). However, only four infected patients were sampled 14 dso in this panel. When combining IgM and IgG results using LFA devices in panel 1, the sensitivity was 82% and 71% for LFA-1 and LFA-2, respectively (Fig. 3).

**Figure 3:**
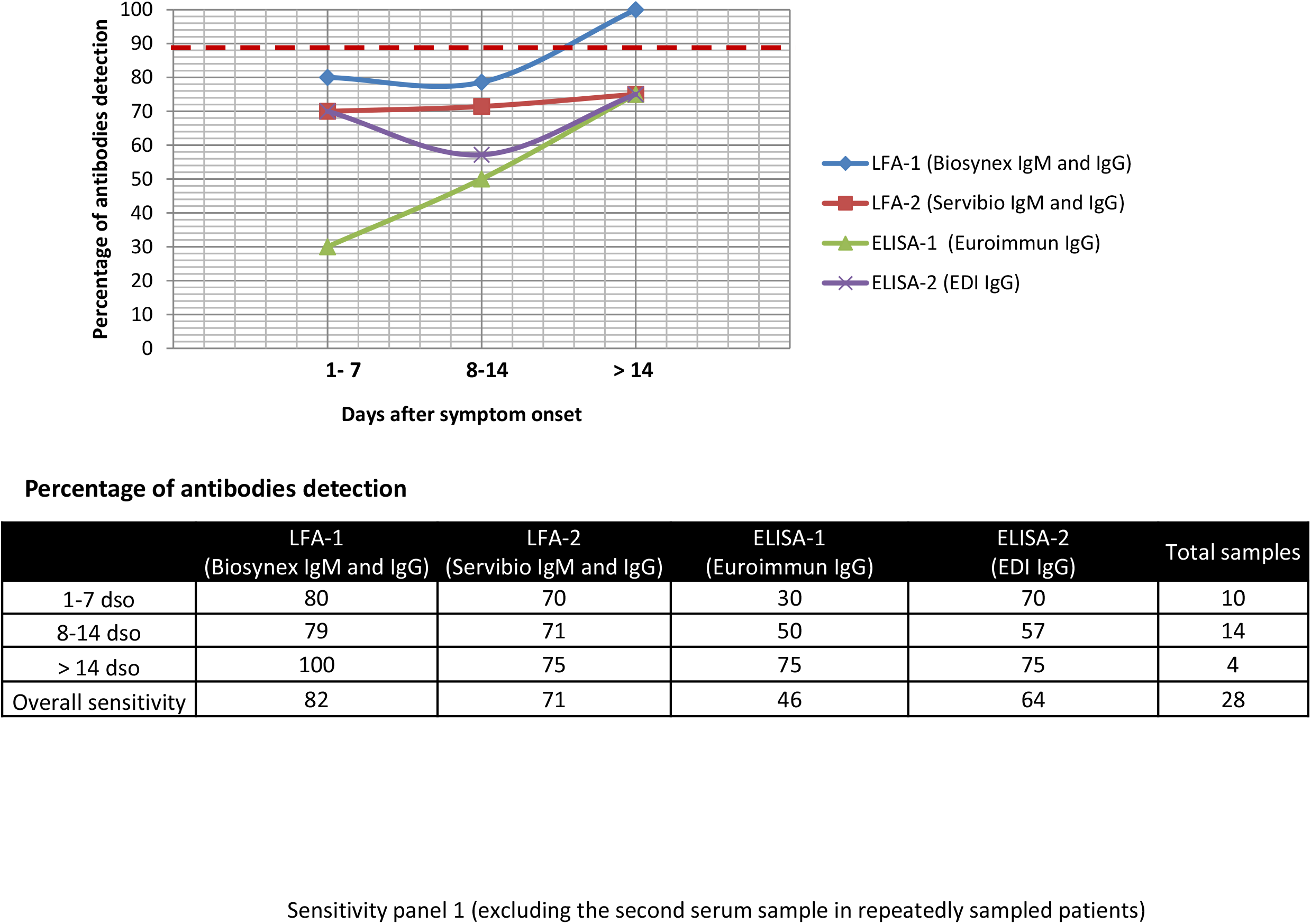
Rates of positivity for virus-specific antibodies measured by LFA (both IgG and IgM) and ELISA (IgG) versus time (in days) after the date of symptom onset in 30 serum samples obtained from COVID-19 hospitalized patients.

In panel 2, the sensitivity for IgM detection was 87% and 29% for LFA-1 and LFA-2, respectively. LFA-1 was more efficient at detecting IgM from 15 to 21 dso in 92% of the cases, whereas the highest percentage of IgM detection for LFA-2 was measured 28 dso, with only 35% of the cases detected (Fig. S5). The percentage of IgG detection ranged from 64% for ELISA-2 to 87% for ELISA-1 (Fig. S6). The optimum rate of IgG detection was observed for all assays after 28 dso (i.e., 68% (ELISA-2), 80% (LFA-2), 88% (LFA-1) and 100% (ELISA-1)). Combining IgM and IgG detection in this panel increased the overall sensitivity to 95% for LFA-1 and to 70% for LFA-2 (Fig. 4).

**Figure 4:**
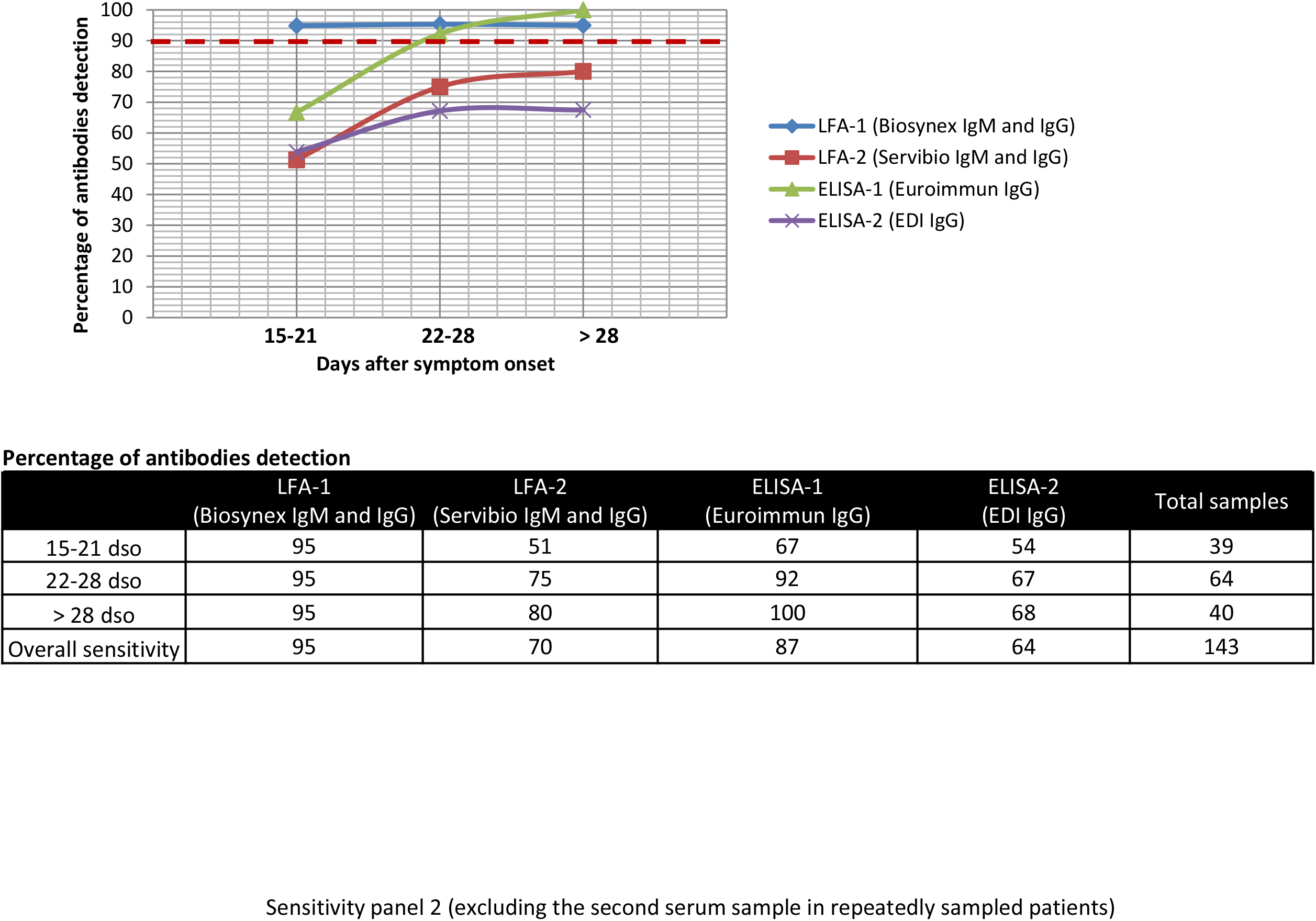
Rates of positivity for virus-specific antibodies measured by LFA (both IgG and IgM) and ELISA (IgG) versus time (in days) after the date of symptom onset in 143 COVID-19 healthcare workers.

IgM clinical specificity ranged from 88% (LFA-2) to 98% (LFA-1). IgG specificity was 98% for LFA-1, whereas it reached only 83% for LFA-2, corresponding to 7/40 false-positive results with a weak intensity score of 1 to 2. ELISA-1 and ELISA-2 showed specificity values for IgG of 98% and 90%, respectively. ELISA-1 showed a specificity of 88% for IgA, and ELISA-2 showed a specificity of 98% for IgM (Table S2).

### Relative performances of serological tools for SARS-CoV-2 (Panels 1 and 2)

The relative performance of evaluated assays was assessed on both panels 1 and 2 (not in panel 3). The overall agreement among the four assays was 79% (Fleiss’ kappa: 0.57; 95% confidence interval [CI]: 0.51-0.62). When comparing the two LFAs, the kappa agreement statistic was 0.50 (95% CI: 0.392-0.615) for IgG and 0.11 (95% CI: 0.014-0.199) for IgM. Between the two IgG ELISAs, the kappa value reached 0.54 (95% CI: 0.433-0.654). High variability in signal intensities was observed among the tested assays (Fig. 5A). Bland-Altman analysis of the IgG ratio measured by ELISA-1 and ELISA-2 defined a 95% limit of agreement of 4.93 (S/CO), showing a good correlation between the two IgG ELISAs with ratios of at least 2 S/CO (Fig. 5B).

**Figure 5:**
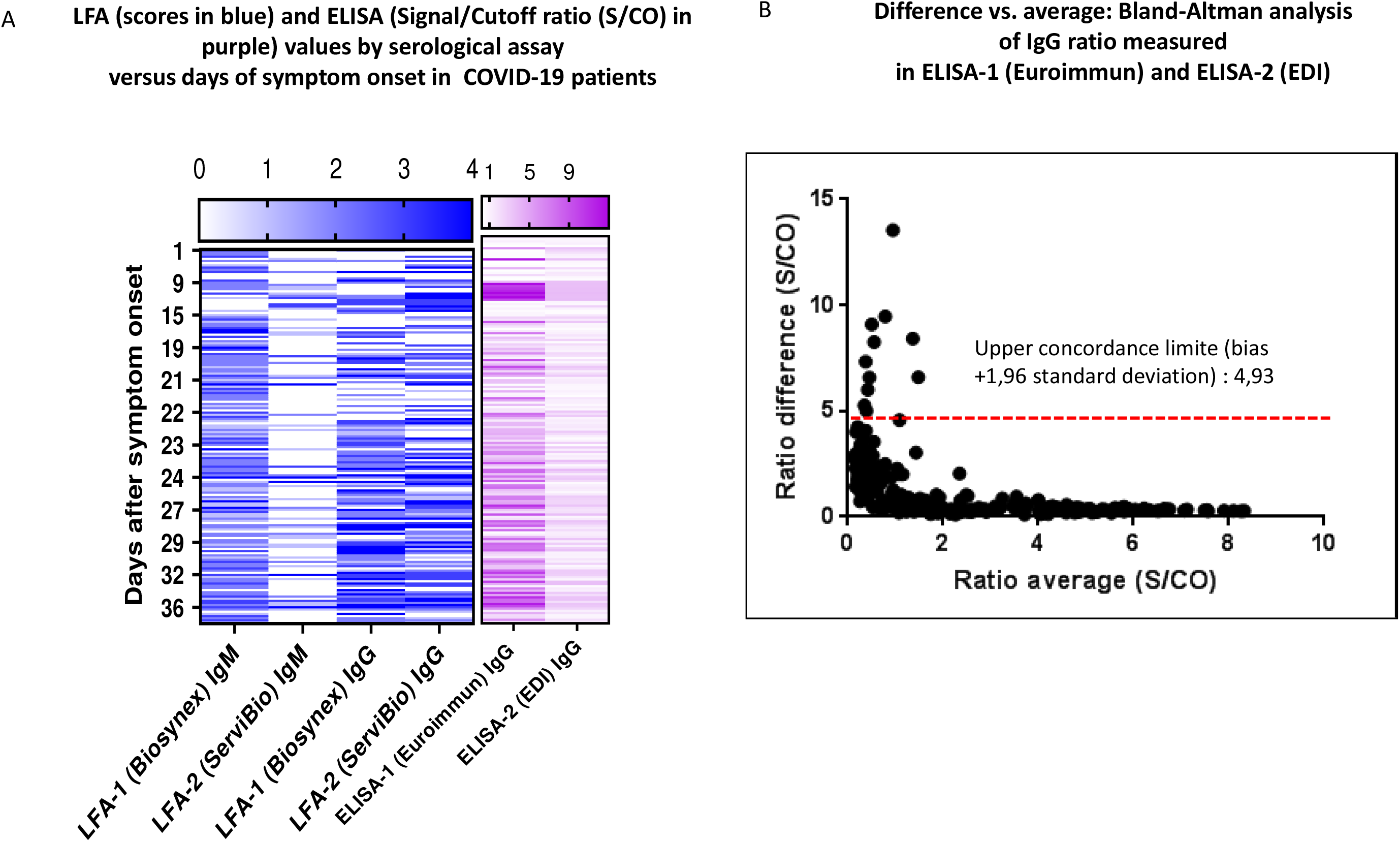
Relative performance of serological tools for the detection of SARS-CoV-2 (panels 1 and 2). (A) LFA (scores in blue) and ELISA (signal/cutoff ratio (S/CO) in purple) values by serological assay versus days after symptom onset in 171 COVID-19 patients. (B) Bland-Altman analysis (difference versus average) of the IgG ratio measured by ELISA-1 (Euroimmun) and ELISA-2 (EDI).

### Time to IgM and IgA antibody onset

Fifty early serum samples (panel 3) of COVID-19 patients were tested with ELISA IgA (Euroimmun) and ELISA-IgM (EDI) assays as well as with both LFA devices. The IgM detection rate ranged from 34% (ELISA-IgM EDI) to 48% (LFA-1), whereas IgA was detected in 40% of samples. The optimum rate of detection for IgM and IgA was observed between 9 and 14 dso (82% for LFA-1 IgM and 71% for IgA ELISA) (Fig. 6). We further analyzed the delay of antibody onset in this panel according to the hospitalization unit. When considering samples positive in at least two of the four assays, we observed a trend towards an earlier detection of antibodies in patients admitted to the ICU than in those with milder disease, but the specimen numbers per time interval were low (Fig. 7).

**Figure 6:**
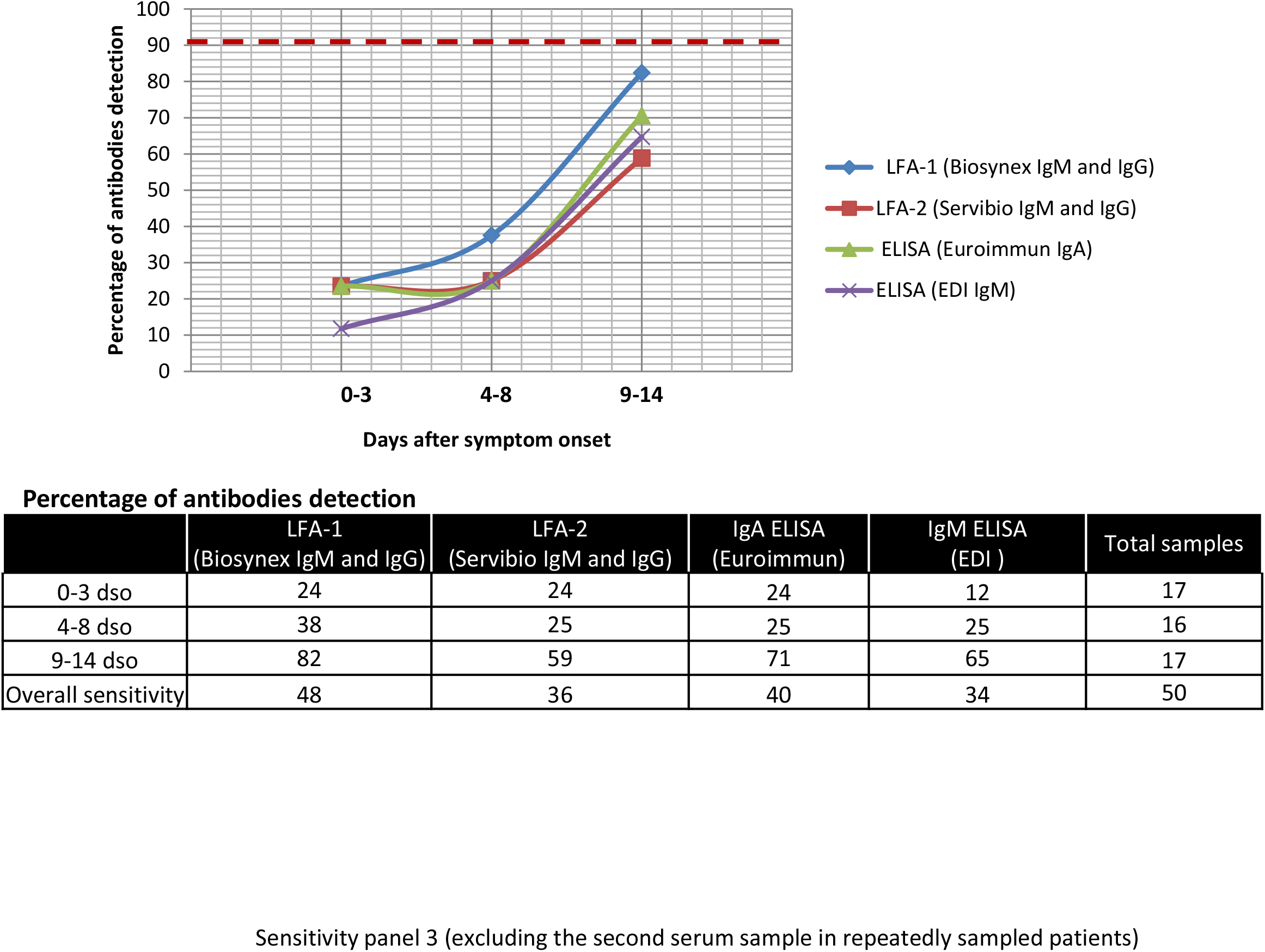
Rates of positivity for virus-specific antibodies measured by LFA (combining IgG and IgM), ELISA (IgA), and ELISA (IgM) versus days after symptom onset in 50 COVID-19 hospitalized patients (panel 3).

**Figure 7:**
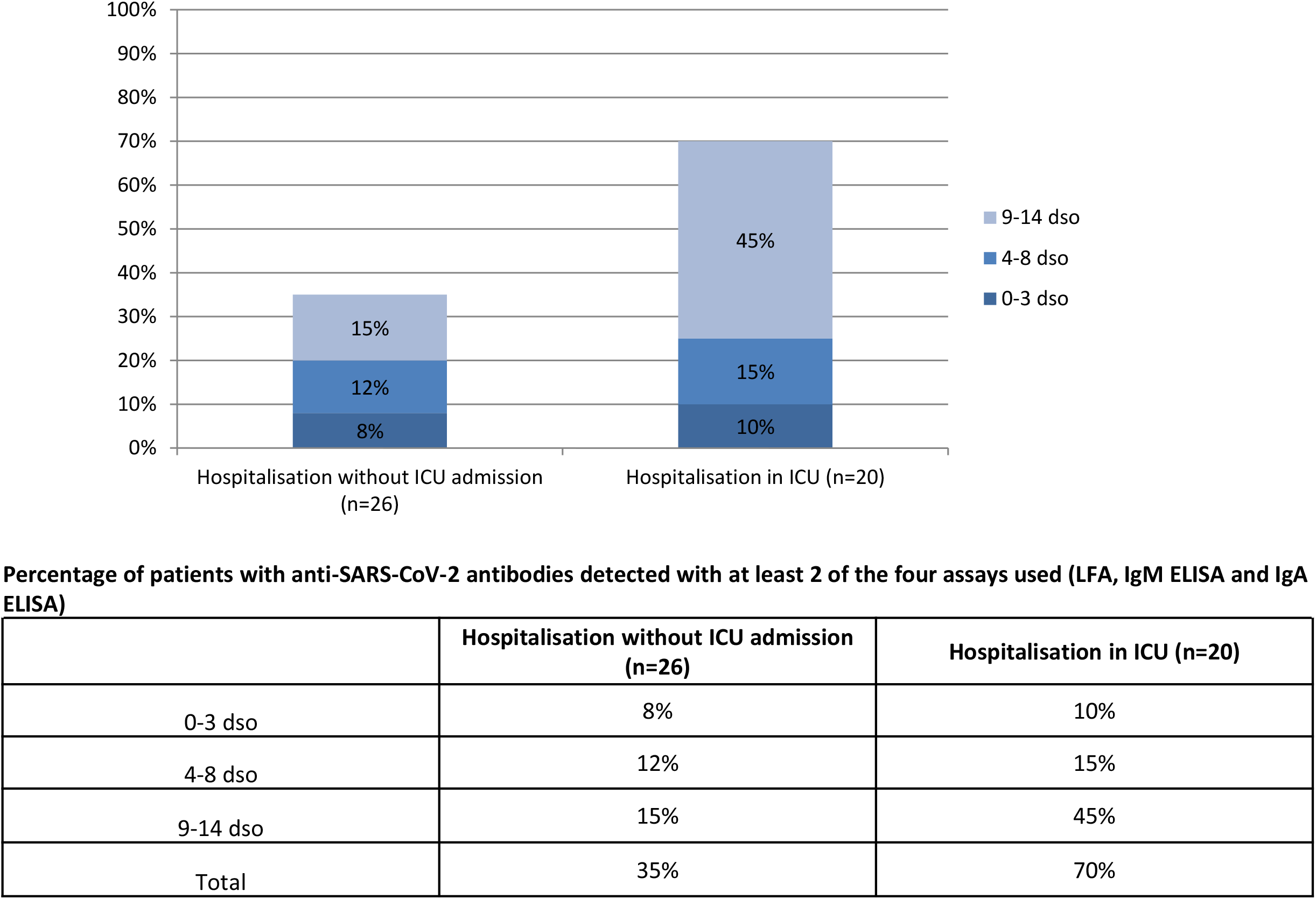
Rates of positivity for virus-specific antibodies according to the need for intensive care unit (ICU) admission.

### Cross-reactivity between SARS-CoV-2 and other Human Coronaviruses

Analytical specificity reached 89% for IgM and 100% for IgG for both LFAs. LFA-1 cross-reacted with the two serum samples containing rheumatoid factor (IgM band intensity scored from 1 to 3). Both LFA assays cross-reacted with seasonal human coronaviruses (HCoV-HKU1/NL63, 229E and OC43) with IgM band intensities scored from 1 to 2 (Table S3). The analytical specificity was 96% for both IgG ELISA devices and reached 93% for IgA ELISA (Euroimmun) and 100% for IgM ELISA (EDI). Both IgG ELISAs cross-reacted with a different seasonal human coronavirus (HCoV-HKU1 for ELISA-2 (EDI) and HCoV-NL63 for ELISA-1 (Euroimmun)).

## DISCUSSION

In our study, we evaluated test performance for two LFAs (i.e., Biosynex (LFA-1) and Servibio (LFA-2)) and two ELISA kits (i.e., ELISA-1 Euroimmun IgA and IgG and ELISA-2 EDI IgM and IgG). We found a good clinical specificity of 98% for LFA-1 (Biosynex IgM and IgG), ELISA-1 (IgG) and ELISA-2 (IgM). Except for ELISA-1 IgA and for the IgM test line on both LFA devices, other assays did not cross-react or they poorly cross-reacted.

Clinical sensitivity was first calculated on combined panels 1 and 2 according to days after symptom onset. Considering the 171 serum samples, the majority of patients displayed anti-SARS-CoV-2 antibodies only 15 days after symptom onset. The assays we tested showed variable sensitivities and poor mutual agreement (Fig. 6A). However, only IgG ELISA-1 (Euroimmun) reached more than 90% clinical sensitivity 21 dso (Fig. 2). The observed differences in terms of sensitivity may reflect the material used as an antigenic source for each assay. Among the 4 coronavirus structural proteins, the spike (S) and nucleocapsid (N) proteins are the main immunogens (8). Specifically, antibodies directed against the viral spike protein are expected to appear earlier than those directed against the nucleocapsid protein (3, 9). ELISA-1 (Euroimmun) and LFA-1 (Biosynex) use the recombinant spike protein S1 domain and the receptor binding domain (RBD) as antigenic sources, respectively, whereas ELISA-2 (EDI) and ELISA-2 (Servibio) are based on a recombinant complete nucleocapsid protein. Another major point explaining the variable results is the choice of the population tested.

Since their development and availability, serological tools have been envisaged to meet two different objectives.

The first objective was to obtain a faster diagnosis, improve the detection of acute infection by detecting false-negative patients to decrease workloads to central laboratories and accelerate clinical decision-making (6, 7). We evaluated clinical sensitivity in panel 1, including serum samples from hospitalized patients with COVID-19. Only LFA-1 (Biosynex) reached 100% detection after 14 dso when combining IgM and IgG detection. For the same time point, the other assays (LFA-2, ELISA-1 IgG and ELISA-2 IgG) showed a suboptimal sensitivity of 75%, which moderates the interest in their use in the triage of patients with suspected COVID-19. Moreover, because of possible delays in seroconversion, we suggest that rapid serology tests such as LFA cannot replace RT-PCR but should instead be considered complementary tools to enhance access to the screening of symptomatic and asymptomatic patients at the population level.

We also investigated IgM and IgA detection in a third panel (panel 3) of sera from infected patients sampled early after symptom onset. If some patients developed anti-SARS-CoV-2 antibodies from 1 to 3 dso, most had detectable IgM (65% with EDI IgM ELISA) and IgA (71% with Euroimmun IgA ELISA) only between 9 and 14 days (Fig. 6). In addition, symptom severity may affect the rate of seropositivity. A delayed or absent humoral response against SARS-CoV-2 has been reported in some patients (10) and may result in negative serology results (7; 11). Future studies are required to shed further light on the underlying mechanisms. We observed a trend towards higher seroprevalence, with at least two of the four assays being positive for patients admitted to the ICU compared to those with milder disease.

The second diagnostic application of a SARS-CoV-2 serological diagnostic tool would be to determine population seroprevalence. At this stage of the pandemic, many countries are now preparing the exit from lockdown. Serological tools have an important place in establishing such strategies. Therefore, we evaluated the four assays in panel 2, composed of 143 serum samples from COVID-19 healthcare workers with a diagnosis proven by RT-PCR. Only LFA-1 (Biosynex) when combining IgM and IgG detection and ELISA-1 IgG (Euroimmun) displayed an excellent clinical sensitivity after 21 days from the onset of symptoms in the range of acceptable values defined by the French National Agency of Medicine and Health Products Safety (ANSM) (90-100%) (Fig. 4).

At this stage of the pandemic, there are no data available about the COVID-19 global seroprevalence in our country (or only partial data obtained in a small specific cohort). It would be interesting in light of future prevalence studies to determine and discuss the positive predictive value (PPV) of the LFA and ELISA kits we evaluated.

In this study, we first demonstrated that serological tools cannot replace RT-PCR for acute infection diagnosis, and second, a thorough selection of serological assays for detecting ongoing or past infections is advisable following the lifting of lockdowns. Special attention should be paid to antigenic sources and validation against RT-PCR results. The reading of sample test lines on LFA devices is still subjective regardless of the manufacturers, especially for weak and/or equivocal bands, requiring a double reading of results. This subjectivity makes it difficult to globalize their use with good reproducibility among healthcare workers. Manufacturers should provide some intensity scale to facilitate the interpretation of these assays. We recommend optimizing antibody detection by combining one LFA and one IgG ELISA in cases of weak or equivocal signals on the LFA. Long-term studies are required to investigate antibody persistence.

## Abbreviations

COVID-19: coronavirus disease 2019;
dso: days after symptom onset;
ELISA: enzyme-linked immunosorbent assays;
LFA: lateral flow assays;
RT-PCR: reverse transcription (RT-) polymerase chain reaction (PCR);
SARS-CoV-2: severe acute respiratory syndrome coronavirus 2

## ACKNOWLEDGMENTS

We thank Véronique Sohn, Anne Moncollin, Axelle Grub, Nathalie Durand, and Elise Sühr for their excellent technical assistance. Pr Fafi-Kremer had full access to all of the data in the study and takes responsibility for the integrity of the data and the accuracy of the data analysis. This study was supported by the Strasbourg University Hospital (COVID-HUS study). We declare no conflicts of interest.

